# Using a System’s Equilibrium Behavior to Reduce Its Energy Dissipation in Non-Equilibrium Processes

**DOI:** 10.1101/291989

**Authors:** Sara Tafoya, Steven J. Large, Shixin Liu, Carlos Bustamante, David A. Sivak

## Abstract

Cells must operate far from equilibrium^1^, utilizing and dissipating energy continuously to maintain their organization and to avoid stasis and death. However, they must also avoid unnecessary waste of energy^2^. Recent studies have revealed that molecular machines are extremely efficient thermodynamically when compared to their macroscopic counterparts^3,4^. There are also tantalizing hints of molecular machines conserving energy while operating out of equilibrium^5,6^. However, the principles governing the efficient out-of-equilibrium operation of molecular machines remain a mystery. A theoretical framework has been recently formulated in which a generalized friction coefficient quantifies the energetic efficiency in non-equilibrium processes^7,8^. Moreover, it posits that to minimize energy dissipation, external control should drive the system along the reaction coordinate with a speed inversely proportional to the square root of that friction coefficient. Here, we test and validate the predictions of this theory by probing the non-equilibrium energetic efficiency of a single DNA hairpin subjected to unfolding and refolding protocols using a dual-trap optical tweezers.

---

Reversible heat engines operating infinitely slowly according to the Carnot cycle do not dissipate energy; their energetic efficiency is limited only by the entropy increase of the surroundings associated with the transfer of heat from a hot to a cold reservoir. In contrast, for engines operating irreversibly, the extra non-equilibrium energy cost associated with carrying out a process at a finite rate further reduces their efficiency^9^. This is the case of biological machines that must operate under signaling, transport, and cell cycle time constraints. For instance, F_o_F_1_-ATP synthase, the primary machine responsible for ATP synthesis, can rotate up to ~350 revolutions/s^10^; the bacteriophage φ29 packaging motor internalizes the 19.3 kbp viral genome into a small capsid at rates of 100 bp/sec—faster than the relaxation rate of the confined DNA^6^; and during sporulation, the *B. Subtilis* DNA translocase, SpoiIIIE, transfers two thirds of its 4.2 × 10^6^ bp genome between mother cell and pre-spore in only 15 minutes, i.e., at a transfer rate of nearly 4,000 bp/s^11^. The finite-time operations of these machines necessarily involve energy dissipation—often in the form of extra work—and it is of great interest to understand how they attain their large (over 70%) energetic efficiencies^12,13^.

Recently, a generalized friction coefficient—which can be obtained from equilibrium measurements—was shown to be the parameter that governs the near-equilibrium energy dissipation during a finite-rate process^7^. Here, we test experimentally the predictions of this theoretical framework and its implications for the thermodynamic efficiency of non-equilibrium processes. To this end, we subject single DNA hairpins to mechanical unfolding and refolding using protocols dictated by this theory; we show that these protocols systematically and significantly reduce energy dissipation during the process. DNA hairpins are ideally suited for this test as the magnitude of the friction coefficient can be tuned by changing the molecule’s length, the free energy difference, the free energy barrier, and the transition rates between its folded and unfolded states^14^.

According to this near-equilibrium linear response theory, the excess power dissipated by a system taken from an initial to a final state by varying a control parameter *λ* according to a protocol (time schedule) *Λ*, is proportional to a generalized friction coefficient *ζ*^7^:

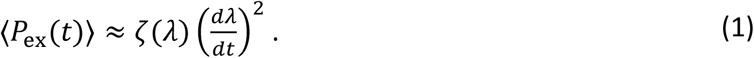

ζ can be computed from the fluctuations *δF*(*t*) = *F*(*t*) — 〈*F*〉 of the force *F*(*t*) via the time integral of the force autocorrelation function 〈δ*F*(0)*δF*(*t*)〉_λ_,

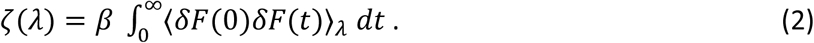

which can be decomposed into

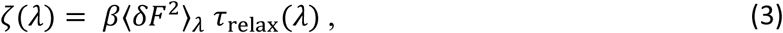

the product of the force variance 〈*δF*^2^〉_*λ*_ and the force relaxation time

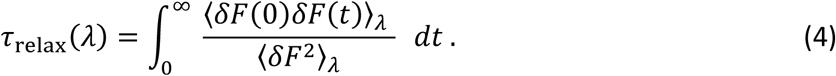

Here, 〈…〉 denotes a non-equilibrium average over system response to a given protocol, whereas 〈…〉_*λ*_ denotes an equilibrium average over system fluctuations at fixed control parameter *λ.*

It can be shown^8^ that near equilibrium, the minimum-dissipation driving protocol, *λ*(*t*)_designed_, proceeds with a velocity proportional to the inverse square root of the friction coefficient *ζ*, *dλ*(*t*)_designed_/*dt* ∝ *ζ* (*λ*)^-1/2^. The proportionality is fixed by the time taken to perform the protocol, so changing the protocol duration corresponds to a global rescaling of all velocities.

To obtain the generalized friction coefficient of the DNA hairpin, we monitored the equilibrium force fluctuations of molecules tethered between two optical traps at various fixed trap separations, *X.* For very small or very large trap separations, the force fluctuates around a single mean value, corresponding to the folded or unfolded conformation, respectively (Fig. 1a); for intermediate trap separations, the force fluctuates between two different values, reflecting the hopping dynamics of the DNA hairpin sampling the folded and unfolded conformations (Fig. 1a). For each separation *X*, we calculated the force autocorrelation function, *〈δF*(*0*)*δF*(*t*)*〉*_*x*_(Fig. 1c); as expected, in the hopping regime the force variance is larger and fluctuations decay more slowly than when an extreme trap separation holds the DNA hairpin in a single conformation.

**Figure 1:**
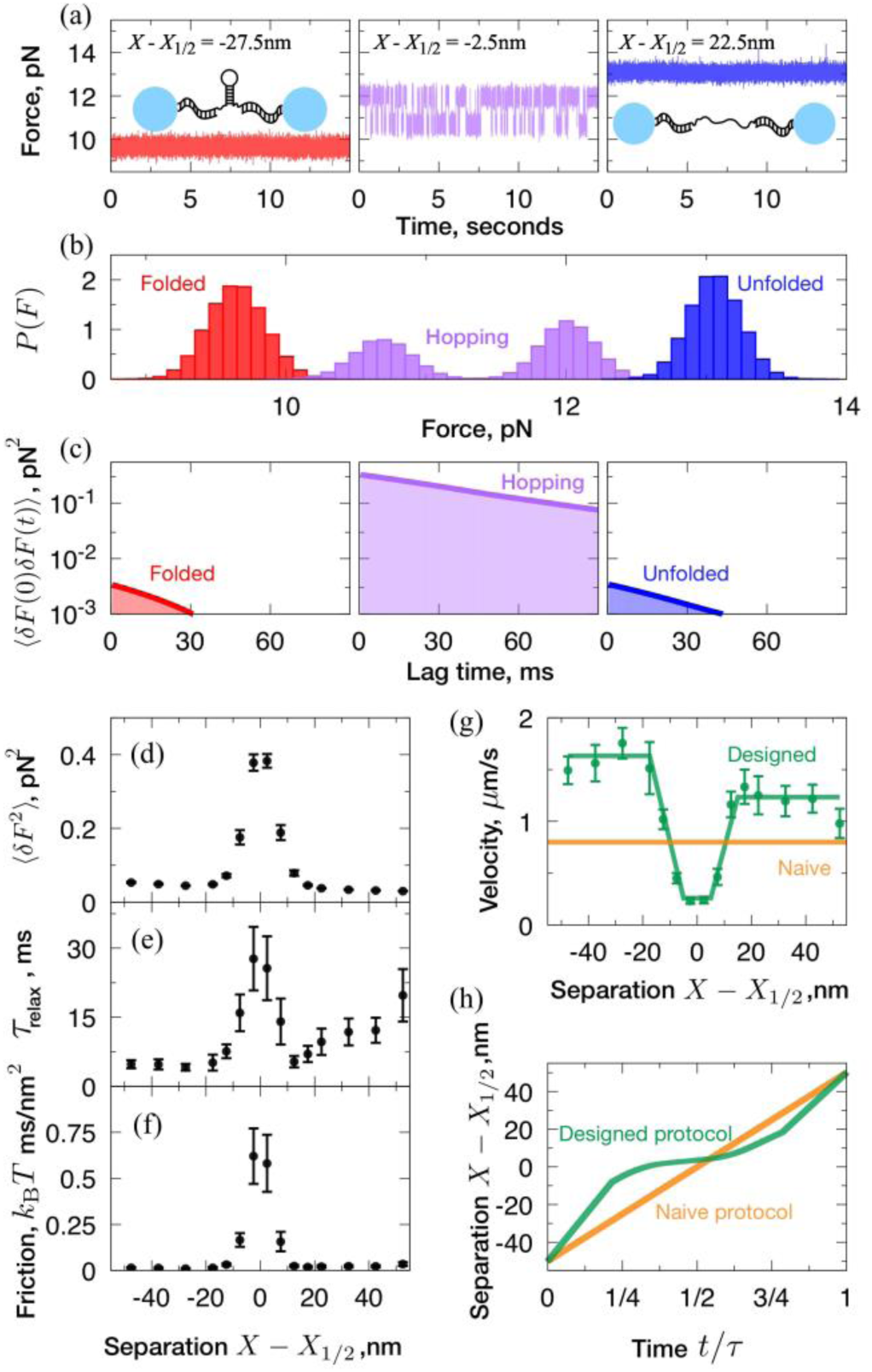
Equilibrium sampling reveals the friction coefficient peaks strongly at the hopping regime. (a) Sample force traces as function of time, for folded hairpin (left, red), hopping hairpin (middle, purple), and unfolded hairpin (right, blue). (b) Equilibrium force distributions and (c) force correlation as a function of lag time, for corresponding fixed optical trap separations. (d) Force variance *〈δF*^2^〉_X_, (e) force relaxation time ^τ^_relax_(*X*) and their product (f) the generalized friction coefficient ζ(*X*), as a function of fixed optical trap separation. (g) For a 0.13-second protocol duration, the designed velocity *dX/dt* ∝ ζ^-1/2^ (green points) with best-fit model (green curve) that minimizes Akaike information criterion ^18^, compared to naive velocity (orange line). (h) Designed and naive velocities scale inversely with protocol duration *τ*, so designed (green) and naive (yellow) protocols are plotted as functions of t/τ

Next, we calculated the force variance (Fig. 1d) and the force relaxation time (Fig. 1e) from the force autocorrelation function. The force variance peaks at an intermediate trap separation, *X_1/2_*, where the hairpin spends roughly equal time between the folded and unfolded conformations. Likewise, the force relaxation time peaks at *X*_1/2_, reflecting that in order to equilibrate, the hairpin must relax across the barrier separating the folded and unfolded states. The generalized friction coefficient—the product of force variance and force relaxation time, equation (3)—also peaks at *X*_1/2_ (Fig. 1f).

As mentioned above, the theory predicts that (near equilibrium) the minimum-dissipation protocol proceeds with a pulling speed—or velocity of the steering trap—that scales as the inverse square root of the friction coefficient^7^: pulling fast at extreme separations where the friction coefficient is small, and slow around *X*_1/2_ where friction peaks. Intuitively, a slow velocity near *X*_1/2_ provides more time for thermal fluctuations to induce the unfolding or folding of the DNA hairpin without additional work input and, therefore, decreases the work required to drive the DNA hairpin between conformations^8^. To ease its implementation, the designed protocol that minimizes dissipation was approximated by a trap velocity profile with a simple piecewise-constant acceleration (Fig. 1g). The resulting designed protocols (Fig. 1h) differ substantially from naive protocols that proceed at constant velocity and that are completed in the same elapsed time. In particular, instantaneous driving velocities varied by a factor of ~6 within a given designed protocol.

Next, we measured force as a function of trap separation, during designed and naive protocols with total durations ranging from 3.7 to 0.13 seconds. These force-separation curves of naive and designed protocols display significant differences in the force at which the DNA hairpins unfold/refold (Fig. 2a). Figure 2b shows the distributions of unfolding force differences, 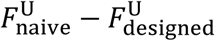, and refolding force differences, 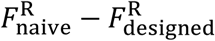, obtained for three different protocol durations. As predicted by theory, on average the DNA hairpin unfolded at lower forces and refolded at higher forces during the designed protocols than during the naive protocols, and the magnitude of the mean force difference is greater for faster protocols (Fig. 2c). These results imply that the designed protocols display lower hysteresis than naive ones (Fig. 3a), a trend that is more prominent in faster protocols where the system is driven farther from equilibrium.

**Figure 2:**
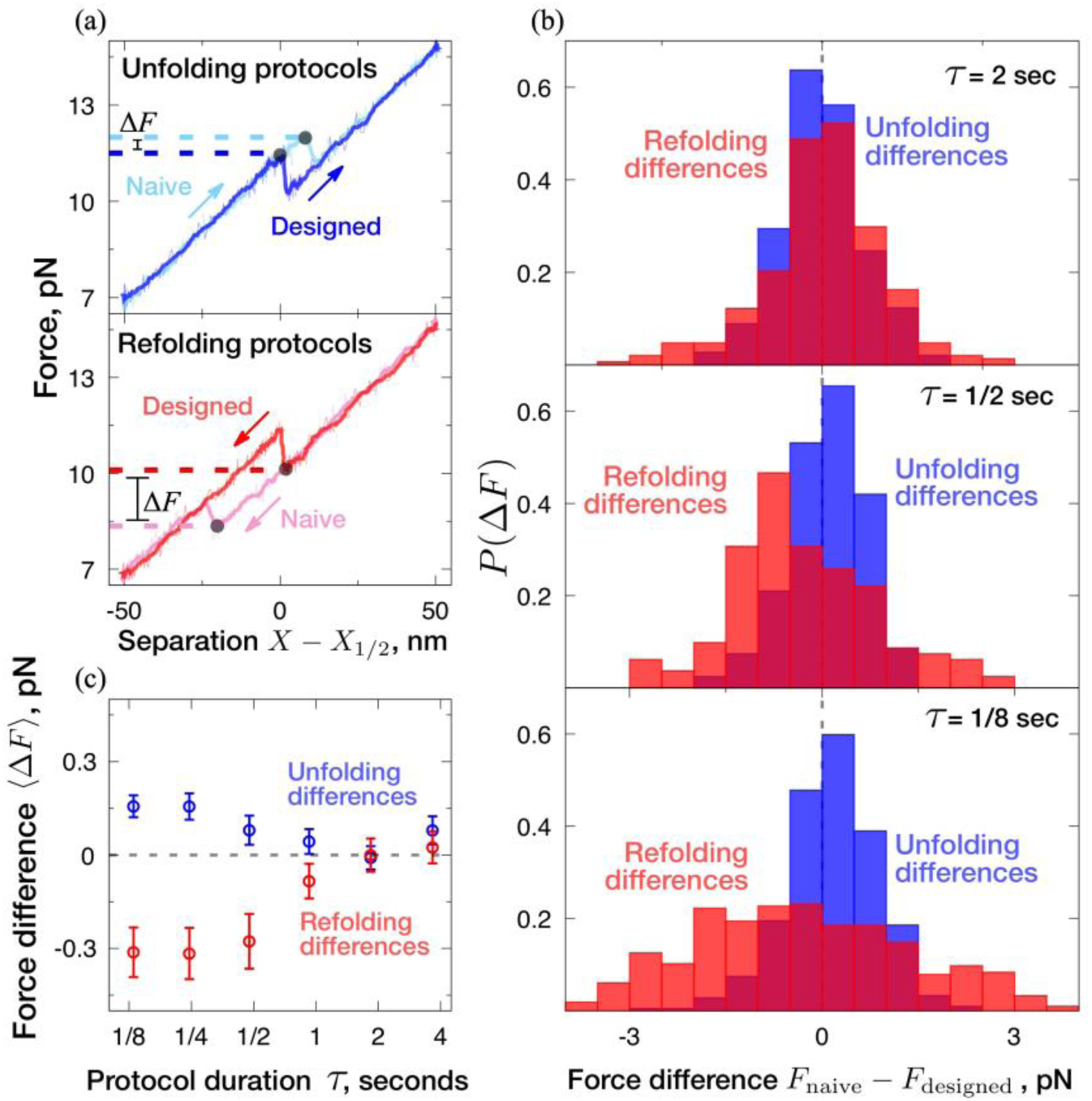
Designed protocols consistently unfold at lower force and refold at higher force. **(a)** Example force-separation curves from a sample molecule for protocol duration *τ* = 0.13 seconds, highlighting the unfolding (top) and refolding (bottom) events (black dots) and the corresponding forces (dashed lines) for designed (dark blue and red) and naive protocols (light blue and pink). The raw data (thin lines) are Savitsky-Golay filtered to obtain a smoothed force-separation curve (thick lines). (b) Distributions of differences *F*_naive_ – *F*_designed_ between naive and designed unfolding (blue) and refolding (red) forces. (c) Mean and standard error for unfolding and refolding force differences, as function of protocol duration. On average, the designed protocol unfolds at a lower force, and refolds as a higher force, than does the corresponding naive protocol.

**Figure 3:**
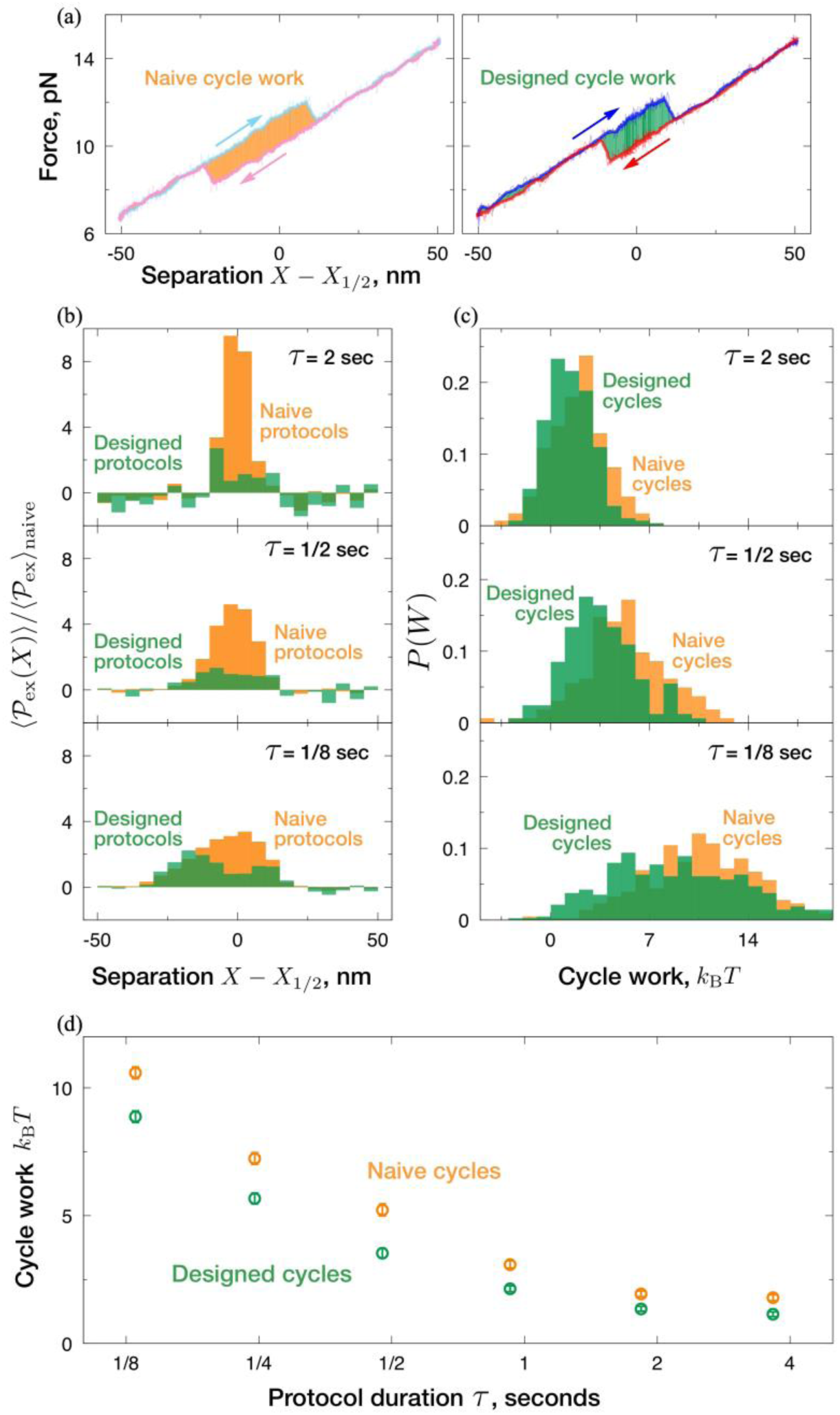
Designed protocols consistently require less work than corresponding naive protocols. (a) Example force-separation curves showing the cycle work *W*^U^+*W*^R^ for naive (left, orange) and designed (right, green) protocols. The raw force-separation curve (thin) is smoothed by a Savitsky-Golay filter (thick). (b) Excess power 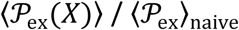 normalized by average naive excess power, as a function of trap separation for naive (yellow) and designed (green) protocols. (c) Distributions of cycle work *W*^U^*+W*^R^, for naive (yellow) and designed (green) protocols, for protocols ranging from slow (top) to fast (bottom). (d) Mean cycle work 〈*W*^U^+*W*^R^〉 during naive (green) and designed (orange) protocols as a function of protocol duration.

According to equation (1), when driving a system at a constant velocity, more work is dissipated at trap separations where the friction coefficient is larger. Consistently, the constant-velocity protocols produce higher dissipation around *X*_1/2_ for all duration (Fig. 3b); by contrast, designed protocols show a substantially flatter dissipation profile across different trap separations (Fig. 3b) and, overall, they induce consistently less dissipation during an unfolding-refolding cycle than do naive protocols (Fig. 3c and 3d). The mean and variance of the distribution of cycle work (*W*^U^+*W*^R^) for both protocol types are higher at shorter protocol duration (Fig. 3c), consistent with higher hysteresis. Finally, the shorter designed protocols save even more work relative to their naive counterparts than faster protocols do (Fig. 3d).

The data presented here correspond to a DNA hairpin that allowed relatively rapid folded-unfolded equilibration, such that transitions to the folded or unfolded conformations occurred even for 0.13-second protocols. This feature allowed us to interrogate the hairpin’s non-equilibrium response over a broad range of protocol durations. In Supplementary Information we show that these results hold also for a different DNA hairpin sequence with significantly (~100x) slower equilibration.

In summary, we have sampled the equilibrium force fluctuations in DNA hairpins, displaying the dynamics of a two-state system (Fig. 1). We showed that the generalized friction coefficient—determined from such equilibrium fluctuations—can be used to design driving schedules (Fig. 1g) that significantly reduce the excess work dissipated compared to constant-velocity schedules (naïve protocols) completed in the same total time (Fig. 3d). This result held for protocol durations that vary by a factor of ~30 (Fig. 3d).

These results have immediate applications in the streamlining of single-molecule experiments and steered molecular dynamics simulations^15^. For instance, to infer the free-energy difference in a given process (such as protein unfolding), the farther the system is from equilibrium during experiment or simulation, the noisier the information obtained about equilibrium properties^16^. Therefore, by sampling the equilibrium fluctuations of a bio-molecular process it should be possible to estimate the generalized friction coefficient across the control parameter landscape, and then craft non-equilibrium protocols that dissipate significantly less energy and achieve significantly greater precision and accuracy for a given number of experimental realizations.

The agreement of theory^7^ and our experiments suggests extensions to more complex contexts. For example, the rotary motor F_1_-ATP synthase is known to be a remarkably efficient machine^12^; such molecular motors may have evolved to slow down their operation in regions of their control parameter space corresponding to high values of the friction coefficient, as a way to harness fluctuations from the thermal bath, thus improving their operation efficiency. Using the above procedure, it would be possible to extract the equilibrium torque fluctuations of the rotary subunit of F_1_-ATP synthase to determine its friction coefficient as a function of the angular control parameter. One could then estimate the minimum-dissipation protocol to drive this subunit. The excess work required by the designed protocol at a given speed provides an approximate lower bound on the energetic costs associated with driving the system out of equilibrium and, thus, sets the scale or metric for judging the non-equilibrium performance of a molecular machine that must turnover on that timescale. Comparing this bound with the empirical efficiency of this motor would quantify how closely the motor’s natural operation has evolved towards minimum non-equilibrium dissipation.

We have seen here that the linear-response theory provides a useful qualitative guide to design protocols that systematically require less work than naive ones. Moreover, this theoretical framework naturally generalizes to stochastic protocols^17^, promising even closer matching to autonomous machines driven by fluctuating forces. Insights from this framework should provide a deeper understanding of the non-equilibrium energetic efficiency of biomolecular machines and ultimately guide the design of efficient synthetic nano-machines.

## Acknowledgements

This work is supported in part by the U.S. Department of Energy under contract number DE-AC02-05CH11231 (to C.B.), the Howard Hughes Medical Institute (to C.B.), the UC-Mexus graduate fellowship (to S.T.), Natural Sciences and Engineering Research Council of Canada (NSERC) CGS Masters and Doctoral fellowships (to S.J.L.), an NSERC Discovery Grant (to D.A.S.), the Faculty of Science, Simon Fraser University through the President’s Research Start-up Grant (to D.A.S.), and a Tier-II Canada Research Chair (to D.A.S.). The authors thank Nancy Forde, John Bechhoefer, Aidan Brown, and Kamdin Mirsanaye (SFU Physics), Michael Woodside (University of Alberta Physics), and Ronen Gabizon and Antony Lee (UC Berkeley) for useful discussions.

## Author contributions

S.T., S.J.L., S.L., C.B., and D.A.S. designed research; S.T. performed research; S.J.L. and D.A.S. analyzed data; and S.T., S.J.L., S.L., C.B., and D.A.S. wrote the paper. ‡ S.T. and S.J.L. contributed equally to this work. *To whom correspondence should be addressed.

## Materials and Methods

### Basic optical trap setup

High-resolution force-separation measurements were conducted on a dual-trap instrument using a solid-state 1064-nm laser, as described previously ^1^. Traps were calibrated as previously described ^2^. DNA tethers were formed between a 0.90-μm-diameter streptavidin-coated bead and a 1-μm-diameter anti-digoxigenin-coated bead (Spherotech) held in separate optical traps. An oxygen scavenging system (100 μg ml^-1^ glucose oxidase, 5 mg ml^-1^ dextrose (Sigma-Aldrich) and 20 μg ml^-1^ catalase, Calbiochem) was included in the buffer to prevent the formation of reactive singlet oxygen, thus increasing the lifetime of the DNA tethers.

## DNA molecules

Hairpin DNA sequences were selected to display hopping dynamics such that determining *X*_1/2_ was accessible experimentally — very fast hopping dynamics were difficult to distinguish from noise, and very slow dynamics required long periods of data acquisition and laser exposure prior to pulling experiments. Minimizing laser exposure avoids molecule photo-damage. All data in Supplementary Information is from sequence 1, GAGTCCTGGATCCTGTTTTTTTTCAGGATCCAGGACTC, which was previously characterized and exhibited appropriate hopping dynamics *t*_1/2_ ≈ 0.24 s ^3^). All data in the main text is from sequence 2, TACCTGATCAGGTGCTTTTTTTTGCACCTGATCAGGTA, the result of modifying sequence 1 to increase GC-content at the loop neck. This change in sequence is expected to facilitate nucleation of the native conformation and to avoid molecule mis-folding ^4^.

Bead size variation, small differences in chemical attachments, and non-specific interactions with the bead surface can lead to molecule-to-molecule variation. We minimized the contribution of trap distance variation by subtracting the value of *X*_1/2_ in all cases. However, other unaccounted sources, such as error in stiffness calibration (most commercially available beads have root-mean-square variations in radius of 3–6%, so the error in stiffness calibration is ~4% assuming a 4% error in reported bead size using individual calibration measurements for each bead pair) and natural variation in the molecules’ persistence length (the standard deviation in persistence length measurement can be as high as ~17% ^5^), also contributed to molecule-to-molecule variation in unfolding/refolding trajectories.

## Equilibrium sampling

Each of 20 molecules is initially probed to find *X*_1/2_: the distance between the traps is increased gradually until the residence time at folded and unfolded conformations is ~50%. Upon identification of *X*_1/2_, a systematic error of −2.5 nm was introduced in the absolute distance between the two traps. This error was introduced as a small difference of a few mV between the instruction given by the computer and the actual analog number instructed to the steering mirror of the trap. This problem is not present when measuring changes in separation, because in calculating relative distances the offset is canceled. We theoretically estimated the error introduced in a designed protocol offset by this amount, and found that such error should lead to a cycle work overestimate of ~6%.

For each molecule, each separation is sampled for 30 seconds, in order from smallest (X_1/2_ – 50 nm) to largest separation (*X*_1/2_ + 50 nm), at 10 nm spacing far from *X*_1/2_ and 5 nm spacing near *X*_1/2_ to more precisely resolve the friction variation at the hopping regime. Changes in separation are instructed to be performed instantaneously but are limited by the response of the mirror controlling the steering trap (~2 ms).

Equilibrium force fluctuations at each of several fixed separations were measured independently in each of 20 different molecules. From these fluctuations the generalized friction coefficient was estimated using equation (2). At each separation, we jackknife resampled from the set of 20 friction estimates to calculate the mean generalized friction and standard error ^6^.

We fit several piecewise-constant-acceleration profiles of protocol velocity to the minimum-dissipation one *dλ_esigned_*/*dt* ∝ [ζ(λ)]^-1/2^ predicted from the empirically determined generalized ζ(*λ*). Each model velocity profile has constant velocity (zero acceleration) far away from *X*_1/2_ and in the immediate vicinity of *X*_1/2_. Constant-acceleration regions interpolate between these constant-velocity regions. The model parameters are the region boundaries and the constant velocities. Different model velocity profiles impose different symmetries, such as inversion symmetry about *X*_1/2_, thus reducing the number of free parameters. We used the velocity profile (Fig. 1) that minimized the Akaike Information Criterion ^7^, a measure of a model’s balance between accuracy and complexity.

## Naive and designed protocols

We estimate the work *W* during a trajectory of forces *F*_*i*_ and separations *X*_*i*_ at *n* discrete time points, by numerical integration

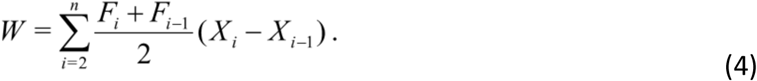

There are 14, 9, 8, 8, 10, and 9 separate molecules sampled with 888 (444), 590 (295), 396 (198), 590 (295), 592 (296) and 472 (236) individual realizations (full cycles) of protocols for durations of 0.13, 0.24, 0.48, 0.93, 1.8 and 3.7 seconds, respectively. The cycle work (hysteresis) 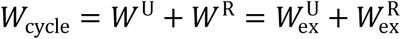 sums the forward and reverse realizations of a protocol at a given speed within protocols taken from the same molecule. By canceling the equilibrium free energy changes during the unfolding and refolding trajectories, this gives the sum of the excess work in each direction.

We investigate 6 different protocol durations, ranging from 0.13 to 3.7 seconds. For each protocol duration, we calculate the work along ~1200 individual realizations, ~300 of each of the 4 protocol types: designed or naive and unfolding or refolding.

To estimate the unfolding (refolding) force in a given force-separation curve, we first smooth the force trace using a second-order Savitsky-Golay filter with window width ~0.4 ms. We report the unfolding (refolding) force as the maximum (minimum) force before the final unfolding (refolding) event takes place. We control for intermolecular variation by analyzing the difference between unfolding/refolding forces along naive and designed protocols for a given molecule, instead of raw unfolding/refolding forces.

The excess power in a protocol interval (Fig. 3b) is calculated by adding the total unfolding work in an interval Δ*X* to the total refolding work in the same interval and dividing by the time taken for the protocol to traverse that separation interval. Finally, the power in each interval is normalized by the average naive excess power (averaged over the entire protocol).

## Supplementary Information

Here we present data from the DNA hairpin sequence presented in the main text (left column in figures), alongside corresponding data from an alternative DNA hairpin sequence (right column).

Fig. S1 shows that (a) the two hairpin constructs have similar force variance as a function of separation, (b) the alternative hairpin sequence has a much greater relaxation time, and thus (c) the alternative hairpin sequence has a much greater generalized friction coefficient. Despite these differences, since the designed protocol only depends on relative variation of the generalized friction coefficient, (d) the designed control parameter velocity and (e) the designed protocol are broadly similar for the two different hairpin constructs.

Fig.S2 shows that the slow-relaxing construct also consistently requires greater cycle work for a naive protocol than for a designed one, even as protocol duration decreases and each cycle work distribution shifts to higher values. As expected given its much greater friction coefficient, non-equilibrium unfolding and refolding of the slowly relaxing hairpin construct produces means and distributions of cycle work shifted to significantly higher values than seen for the rapidly relaxing hairpin. As for the rapidly relaxing hairpin, across all protocol durations for the slowly relaxing hairpin, the designed protocol systematically requires less cycle work than the naive one.

---

**Figure S1:**
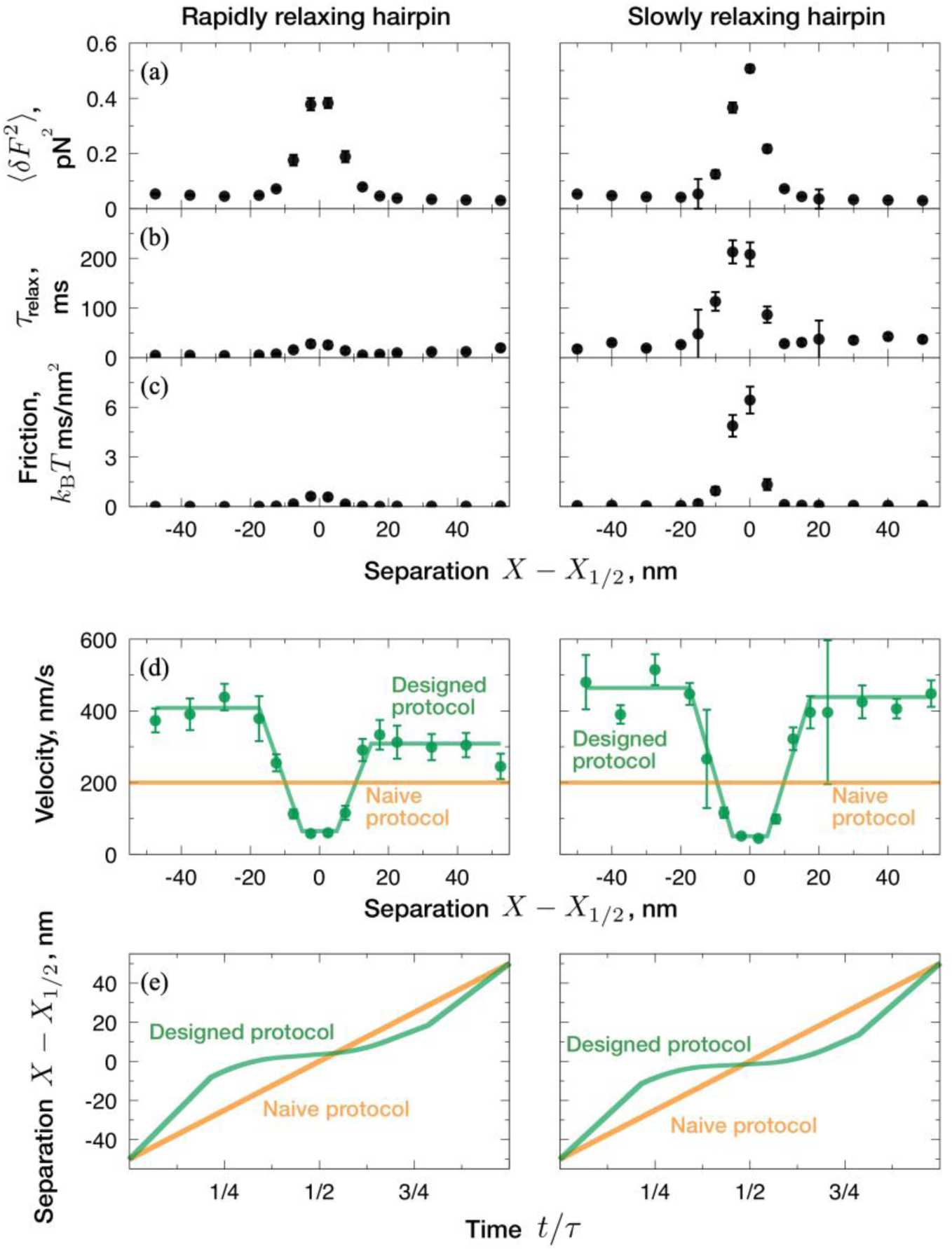
(a) Force variance 〈δ*F*^2^〉, (b) integral relaxation time ^*τ*^_relax_, and (c) generalized friction for two different hairpin sequences. The rapidly relaxing hairpin (left column) is the one presented in the main text, while the slowly relaxing hairpin (right column) has an alternative sequence with friction that is an order of magnitude larger due to increased relaxation time. (d) Implied minimum-dissipation velocity from 20 distinct molecules for each of the rapidly relaxing and slowly relaxing hairpin sequences (dots). (e) Resulting designed protocols (green curves) compared with naive protocols (orange lines).

**Figure S2:**
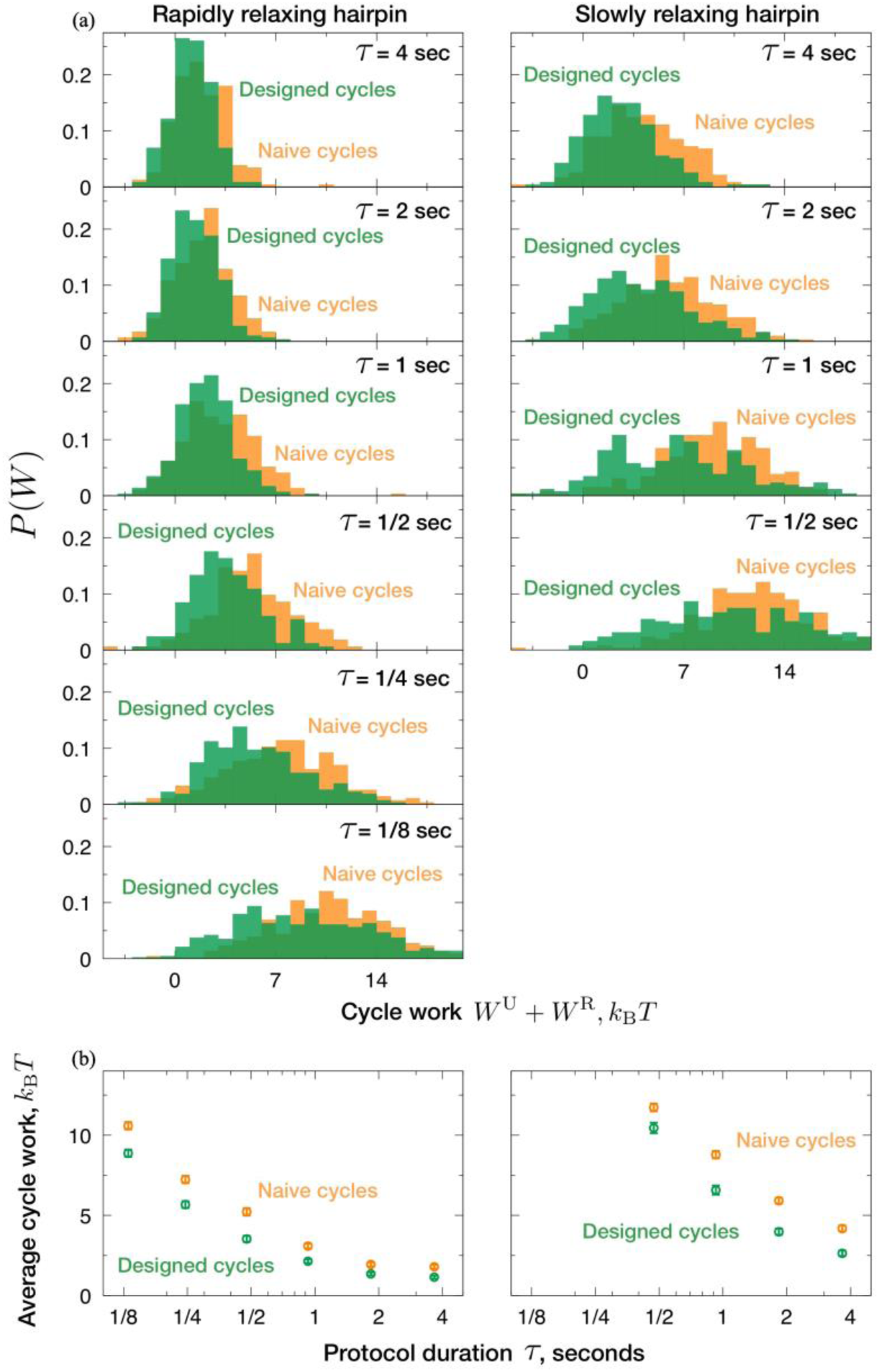
(a) Distributions of cycle work *W*^U^+*W*^R^ for rapidly relaxing (left column) and slowly relaxing (right column) hairpin sequences during naive (orange) and designed (green) protocols, for protocol durations ranging from slow (top) to fast (bottom). Rapidly relaxing histograms are shown for protocol durations of 0.13, 0.24, 0.48, 0.93, 1.8 and 3.7 seconds, with 444, 295, 198, 295, 296, and 236 naive and designed cycles, while slowly relaxing histograms are shown for protocol durations of 3.7, 1.8, 0.93 and 0.48 seconds, with 237, 308, 263, and 257 naive and designed cycles. During the two shortest protocols (0.24 and 0.13 s), the dynamics of the slowly relaxing hairpin did not allow proper refolding, so this data was not analyzed. (b) Mean cycle work 〈*W*^U^*+W*^R^〉 for rapidly (left) and slowly (right) relaxing hairpins during naive (green) and designed (orange) protocols, as a function of protocol duration.

